# SnapHiC-D: a computational pipeline to identify differential chromatin contacts from single cell Hi-C data

**DOI:** 10.1101/2022.08.05.502991

**Authors:** Lindsay Lee, Miao Yu, Xiaoqi Li, Chenxu Zhu, Yanxiao Zhang, Hongyu Yu, Bing Ren, Yun Li, Ming Hu

## Abstract

Single cell Hi-C (scHi-C) has been used to map genome organization in complex tissues. However, computational tools to detect dynamic chromatin contacts from scHi-C datasets in development and through disease pathogenesis are still lacking. Here, we present SnapHiC-D, a computational pipeline to identify differential chromatin contacts (DCCs) between two scHi-C datasets. Compared to methods designed for bulk Hi-C data, SnapHiC-D detects DCCs with high sensitivity and accuracy. We used SnapHiC-D to identify celltype-specific chromatin contacts at 10 kilobase resolution in mouse hippocampal and human prefrontal cortical tissues, and demonstrated that DCCs detected in the cortical and hippocampal cell types are generally correlated with cell-type-specific gene expression patterns and epigenomic features.

## 1. Introduction

The Three-dimensional (3D) architecture of chromosomes in the nucleus plays a key role in the regulation of gene expression^1–4^. Consequently, the disruption of 3D genome structure is often associated with gene dysregulation contributing to a variety of human diseases including cancer^5^. High-throughput chromatin conformation capture technologies (i.e., Hi-C)^6,7^ have been widely used to characterize the spatial organization of chromatin fibers in a broad spectrum of species. However, traditional bulk Hi-C assays require a large volume of input materials, preventing them from capturing cell-type-specific 3D genome organization in complex tissues. In recent years, single cell Hi-C (scHi-C) and related methods^8–13^ have enabled the measurement of chromatin organization in individual cells, facilitating the identification of cell-type-specific 3D genome features directly from complex tissues.

While scHi-C technologies have evolved rapidly, statistical models and computational tools tailored to extract rich information in scHi-C data are still in the early stages of development. Most recent efforts have targeted the characterization of 3D genomic features at single cell resolution, such as A/B compartments and topologically associating domain (TAD)-like structures^14–17^ (more details can be found in the recent review papers^18–20^). However, tools for comparative analysis of scHi-C datasets, for example, identifying differential chromatin contacts (DCCs) between cell types, have yet to be developed.

Several methods have been reported to identify DCCs in bulk Hi-C data from distinct cell types or developmental stages^21–26^. However, no method has been developed for identifying DCC from scHi-C data. In order to detect DCCs between different cell types from scHi-C data, one approach is to first aggregate single cells belonging to the same cell type into pseudo bulk Hi-C data, and then apply existing DCC callers designed for bulk Hi-C. However, such an approach is sub-optimal for scHi-C data for at least two reasons. First, scHi-C data is extremely sparse. It usually requires thousands of cells to achieve sufficient sequencing depth, limiting its use to the most abundant cell types in a tissue sample^16,20^. To address the data sparsity issue associated with scHi-C data, imputing missing contact frequency in each cell becomes a standard preprocessing step for most scHi-C data analysis^18–20^. After imputation, chromatin contact probabilities become a continuous variable taking a value between 0 and 1. Therefore, the negative binomial distribution used in most bulk Hi-C DCC callers^23–25^ cannot fit such continuous data. Second, the sample sizes available for differential analysis are much larger for scHi-C compared to bulk Hi-C data. In bulk Hi-C, most cell types or experimental conditions only consist of a small number of replicates (two or three). The limited number of replicates poses a great challenge in accurately estimating both between and within cell type biological variability. In contrast, scHi-C data typically consist of hundreds of cells of a specific cell type. Treating each cell as an independent unit has the potential to boost the statistical power of detecting DCCs. In sum, existing DCC callers designed for bulk Hi-C data are not optimal for scHi-C data. DCC callers tailored for scHi-C data are of urgent need.

## 2. Results

### 2.1 SnapHiC-D algorithm

We recently developed SnapHiC^16^, the first method to identify chromatin loops at kilobase resolution from scHi-C data. Here, we extend SnapHiC to SnapHiC-D, for comparative analysis between different cell types and to identify DCCs from scHi-C data (see details in **Methods**). Briefly, we first create the 10Kb bin resolution contact matrix for each chromosome in each single cell and then model it as a graph, where each node is a 10Kb bin, and the edge connecting a pair of 10Kb bins is the observed scHi-C contact spanning between these two 10Kb bins. We also add edges for any two consecutive 10Kb bins to make each chromosome a connected undirected graph. Since such graph is extremely sparse due to the limited sequenced depth in scHi-C data, we apply the random walk with restart (RWR) algorithm to impute all the missing edges (i.e., chromatin contact probabilities), as described in the previous studies^14,16^ (**Figure 1A**). Next, we convert the imputed chromatin contact probabilities into Z-scores, for all 10Kb bin pairs with the same 1D genomic distance, in order to normalize the 1D genomic distance effect (**Figure 1B**). SnapHiC-D takes the imputed and normalized chromatin contact frequency (i.e., Z-score) in each cell as the input. For each bin of interest, SnapHiC-D first applies the two-sided two-sample t-test to evaluate the difference in chromatin interaction frequency between two cell types and then reports the test statistics (T) and the corresponding P-values (**Figure 1B**). Next, SnapHiC-D converts the P-values into false discovery rate (FDR) for all 10Kb bin pairs with the same 1D genomic distance. Finally, SnapHiC-D defines bin pairs with |T|>2 and FDR<10% as the DCCs (**Figure 1C**).

**Figure 1.**
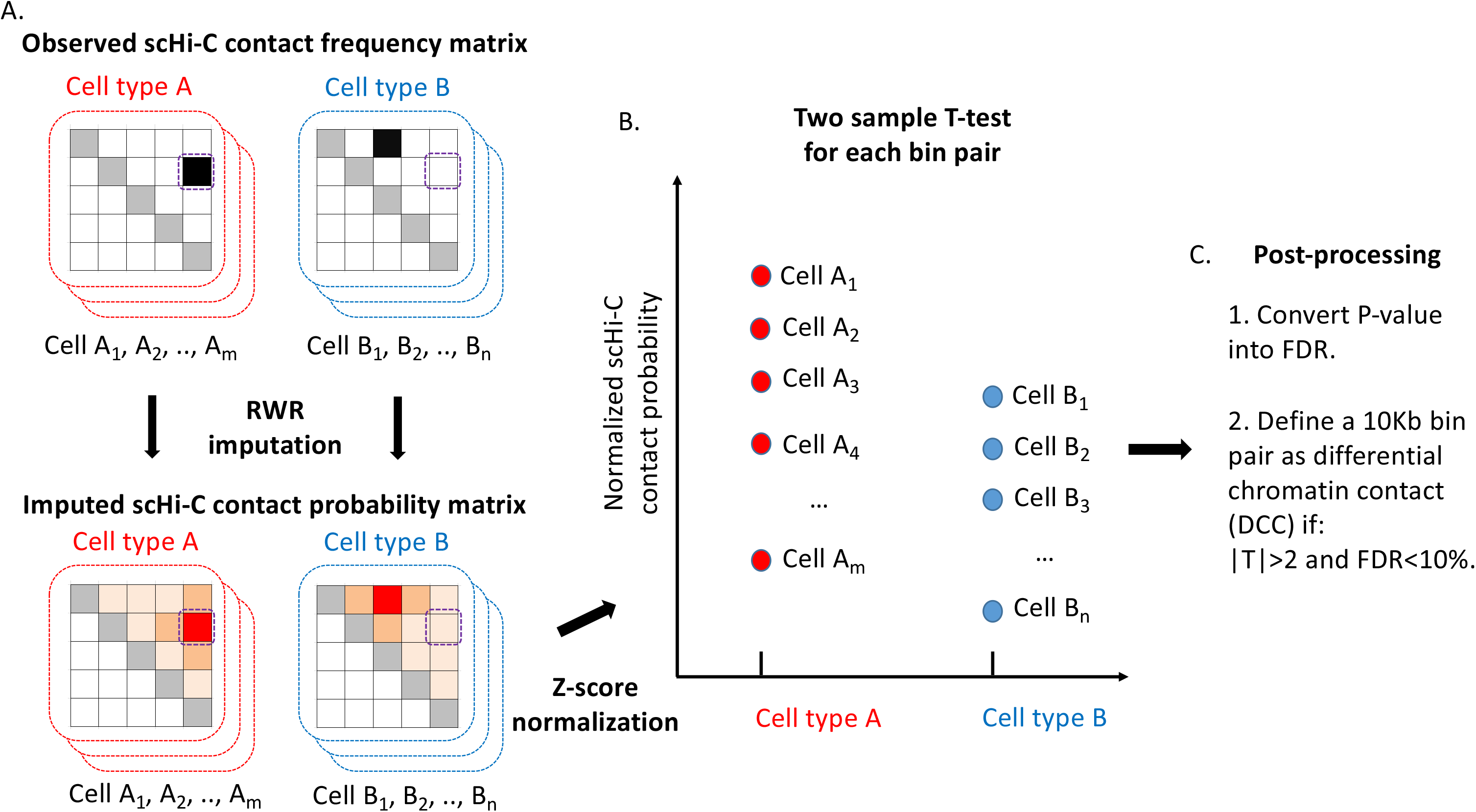
Flowchart of the SnapHiC-D algorithm. **A.** A cartoon illustration of the random walk with restart (RWR) imputation algorithm. Top pattern: the observed scHi-C contact frequency matrix for m and n cells in cell type A and cell type B, respectively. Bottom pattern: the imputed scHi-C contact probability matrix for m and n cells in cell type A and cell type B, respectively. The dashed purple box highlights a bin pair of interest. **B.** A cartoon illustration of the two sample T-test, using the imputed scHi-C contact probability at the bin pair of interest (highlighted by the dashed purple box). **C.** Post-processing step. SnapHiC-D defined a bin pair as DCC if |T|>2 and FDR<10%.

### 2.2 SnapHiC-D controls false positives under the null hypothesis

We first benchmarked the performance of SnapHiC-D under the null hypothesis, which consists of two groups of cells from the same cell type. In this scenario, any identified DCCs should be treated as false positives. We expect that a sensible DCC caller should control false positives. Specifically, we re-analyzed the published scHi-C data from mouse embryonic stem cells (mESCs)^8^, consisting of 742 mESCs with more than 150,000 contacts per cell. We split them equally into two groups A and B and also took the cell cycle stage into consideration (**Table S1**), which could minimize the potential DCCs caused by different cell cycles. These two groups also have a similar number of contacts (the average number of contacts is 442,297 and 442,010 for group A and group B, respectively). We benchmarked the performance of SnapHiC-D against two widely-used DCC callers designed for bulk Hi-C data, diffHiC^23^ and multiHiCcompare^22^. In addition, we used BandNorm to perform scHi-C-specific normalization before applying diffHiC, as suggested by a recent preprint^27^. We did not use the deep learning-based method 3DVI proposed in the same preprint^27^, since BandNorm coupled with diffHiC achieved comparable or superior performance than 3DVI coupled with diffHiC in terms of DCC detection accuracy (see details in Figure S19F in Zheng et al study^27^).

When applying SnapHiC-D to these two groups A and B under the null hypothesis, we observed no change in chromatin interaction frequency at different 1D genomic distances, for all three competing methods (**Figure S1A**, Mean-distance plot [MD plot for short]). SnapHiC-D identified 221 DCCs (|T|>2 and FDR<10%) between group A and group B from the same cell type, among all 528,702 tested bin pairs (thus empirical FDR = 221/528,702 = 0.04%). This empirical FDR suggests that SnapHiC-D rather conservatively controls false positives under the null hypothesis. We further checked the P-value distributions of three alternative methods (**Figure S1B**), and noticed only a slight enrichment of small P-values for SnapHiC-D. In contrast, P-values of BandNorm+diffHiC showed enrichment at the region 0 ~ 0.5 and P-values of multiHiCcompare showed strong enrichment near 1. The abnormal P-value distributions in BandNorm+diffHiC and multiHiCcompare suggested that their statistical models do not fit the sparse scHi-C data, and the FDR calculation is invalid. Therefore, for the remaining part of this paper, we used the nominal P-value <0.05 threshold, rather than FDR to define DCCs for both BandNorm+diffHiC and multiHiCcompare.

### 2.3 SnapHiC-D identifies DCCs with high sensitivity and accuracy

After ensuring SnapHiC-D’s validity in controlling false positives, we next benchmarked the performance of SnapHiC-D against other bulk Hi-C analysis tools in identifying DCC between two different cell types. We generated high coverage scHi-C data from 94 mESCs and 188 mouse neuron progenitor cells (NPCs) (median 1.17 million contacts per cell for mESCs, 1.68 million contacts per cell for NPCs, see details in **Methods**). Noticeably, the number of evaluated bin pairs varies among three methods, where SnapHiC-D evaluated the largest number of bin pairs after imputing missing contacts in each single cell. Specifically, among the 779,772 SnapHiC-D-evaluated bin pairs, SnapHiC-D identified 139,161 DCCs (|T|>2 and FDR<10%, 17.8%), with 55,162 mESC-specific contacts and the rest 83,999 NPC-specific contacts. In contrast, BandNorm+diffHiC identified 80,356 DCCs (P<0.05, 16.7%) among 480,099 BandNorm+diffHiC-evaluated bin pairs, with 40,616 mESC-specific contacts and the rest 39,740 NPC-specific contacts; multiHiCcompare identified 2,881 DCCs (P<0.05, 12.7%) among 22,623 multiHiCcompare-evaluated bin pairs, with 1,192 mESC-specific contacts and the rest 1,689 NPC-specific contacts. Notably, when using the nominal P-value <0.05 threshold, BandNorm+diffHiC and multiHiCompare identified a similar proportion of DCCs compared to SnapHiC-D.

In addition, we checked both the MD plot and P-value distribution of these three methods (**Figure S2**) and found that they are largely consistent with results obtained under the null hypothesis (**Figure S1**). The only noticeable difference is that for multiHiCcompare, NPCs have a higher chromatin contact frequency than mESCs, in particular when the 1D genomic distance is greater than 400Kb. We suspect that such a difference is largely due to NPC (N=188) consisting of more cells than mESC (N=94) and is more pronounced at a large 1D genomic distance (>400Kb).

We further evaluated the sensitivity and accuracy of the 139,161, 80,356 and 2,881 DCCs, identified by SnapHiC-D, BandNorm+diffHiC and multiHiCcompare, respectively. Since no gold standard DCCs between mouse ESCs and NPCs are available, we re-analyzed the deeply sequenced bulk Hi-C data from mESCs and NPCs released in the Bonev et al study^28^, and generated reference DCC lists using the following two approaches (HiCCUPS^7^ and multiHiCcompare^22^). Noticeably, HiCCUPS relies on the local background model to identify chromatin loops, while multiHiCcompare evaluates all bin pairs of interest without the consideration of loop status. Therefore, these two approaches generated different DCC lists. In this work, we treated these reference lists from two complementary methods as the working truth.

First of all, we combined the four replicates of the same cell type together, and applied the HiCCUPS algorithm^7^ to identify 10Kb bin resolution chromatin loops from the combined data. HiCCUPS identified 8,191 and 8,458 loops from mESCs and NPCs, respectively. We then defined cell-type-specific loops, if they are at least 20Kb away from any loops in the other cell type. Since SnapHiC-D only evaluated TSS-anchored bin pairs, to make a fair comparison, we further collected TSS-anchored cell-type-specific loops, leading to a final list of 926 mESC-specific TSS-anchored loops and 1,065 NPC-specific TSS-anchored loops, named as “bulk-Hi-C-specific-loops” for short.

Since bulk-Hi-C-specific-loops are generated from highly stringent HiCCUPS loops, we defined a DCC as “testable” if it is within 20Kb of a bulk-Hi-C-specific-loop. A DCC is considered a true positive if its cell-type-specificity matches with the cell-type-specificity of the bulk Hi-C loop, while it is a false positive if its cell-type-specificity is different from the cell-type-specificity of the bulk Hi-C loop. Among all 139,161 SnapHiC-D-identified DCCs, 2,603 DCCs are testable. For 935 mESC-specific testable DCCs, precision and recall are 68.1% and 20.4%, respectively. For the 1,668 NPC-specific testable DCCs, precision and recall are 79.1% and 25.8%, respectively (**Figure 2A, 2B**). In contrast, among all 80,356 BandNorm+diffHiC-identified DCCs, 1,296 DCCs are testable. For the 707 mESC-specific testable DCCs, precision and recall are 39.6% and 21.4%, respectively. For the 589 NPC-specific testable DCCs, precision and recall are 58.3% and 23.7%, respectively (**Figure 2A, 2B**). Finally, among all 2,881 multiHiCcompare-identified DCCs, only 35 DCCs are testable. For the 13 mESC-specific testable DCCs, the precision and recall are 61.5% and 0.8%, respectively. For the 22 NPC-specific testable DCCs, precision and recall are 50.0% and 0.1%, respectively (**Figure 2A, 2B**). These results suggest that SnapHiC-D outperformed BandNorm+diffHiC and multiHiCcompare in both precision and recall.

**Figure 2.**
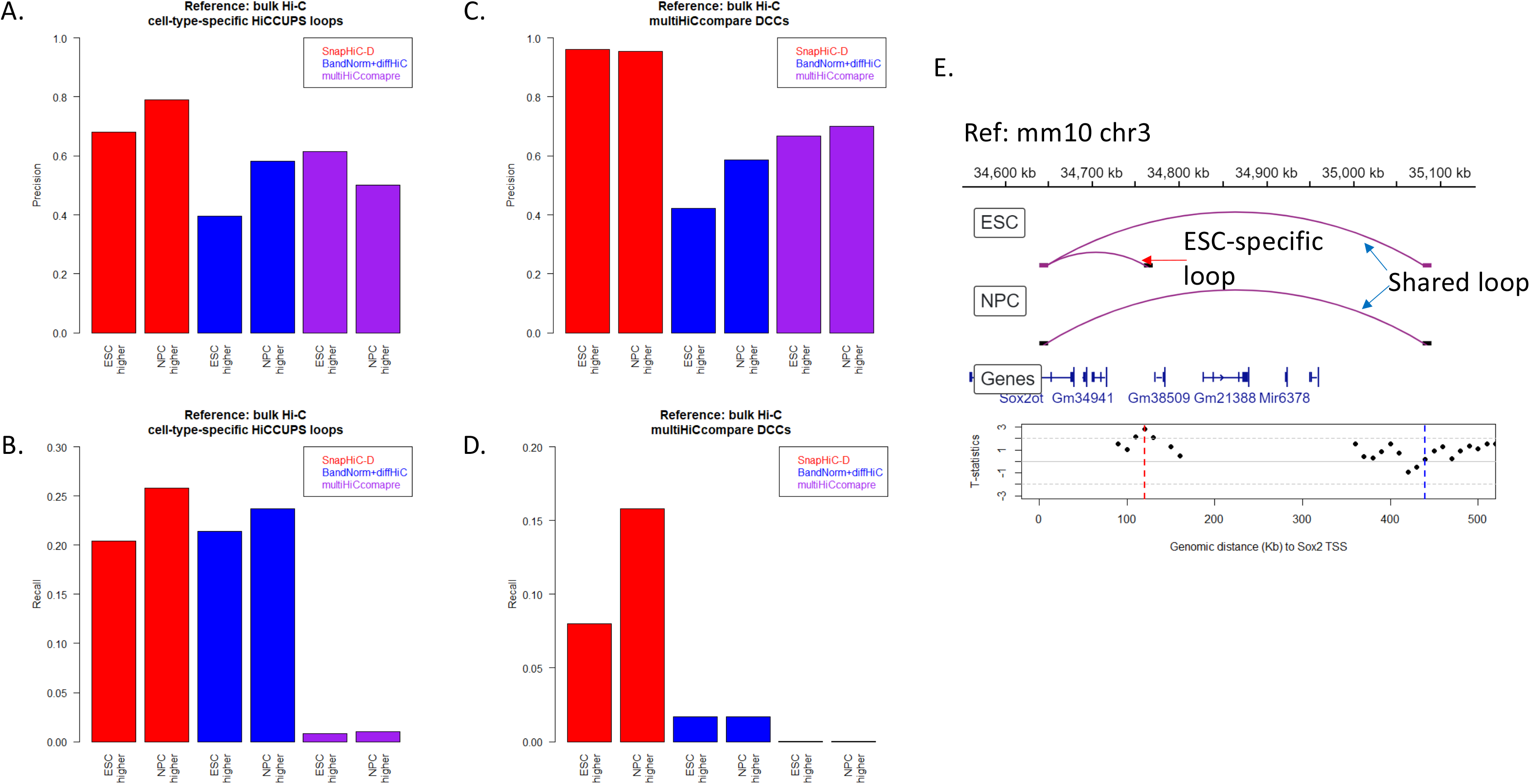
SnapHiC-D identifies DCCs with high sensitivity and accuracy. The precision (**A**) and recall (**B**) of three competing methods using cell-type-specific HiCCUPS loops identified from bulk Hi-C data as the working truth. The precision (**C**) and recall (**D**) of three competing methods using multiHiCcompare DCCs identified from bulk Hi-C data as the working truth. **E.** Top panel: IGV visualization of HiCCUPS loops identified from mESC and NPC bulk Hi-C data, at the *Sox2* TSS. Bottom panel: T-test statistics obtained by SnapHiC-D for tested 10Kb bin pairs anchored at the *Sox2* TSS. The dashed red line and dashed blue line highlighted the location of the mESC-specific loop and shared loop, respectively.

We also benchmarked SnapHiC-D against BandNorm+diffHiC and multiHiCcompare, using DCCs identified by applying multiHiCcompare to bulk Hi-C data as the working truth. Consistent patterns were observed (**Figure 2C, 2D**), similarly supporting that SnapHiC-D outperformed BandNorm+diffHiC and multiHiCcompare in both precision and recall (**Supplementary Material Section 1**).

In addition to the genome-wide comparison, we further checked DCCs at the *Sox2* locus. Using bulk Hi-C data^28^, HiCCUPS identified a ~120Kb mESC-specific loop linking *Sox2* TSS to an mESC-specific superenhancer^29^, and a ~440Kb mESC/NPC-shared loop (**Figure 2E** top panel). We further plotted the T-test statistics, for all testable 10Kb bin pairs anchored at the *Sox2* TSS (**Figure 2E** bottom panel). For the mESC-specific loop, the average normalized contact frequency is 0.94 and 0.17 for mESC and NPC, respectively. The T-test statistic is 2.84 and FDR is 0.06, suggesting that this is an mESC-specific DCC. In contrast, for the mESC/NPC-shared loop, the average normalized contact frequency is 1.93 and 1.81 for mESC and NPC, respectively. The T-test statistic is 0.20 and FDR is 0.91, suggesting that this is not a DCC. As a comparison, neither BandNorm+diffHiC nor multiHiCcompare can detect the mESC-specific loop between the *Sox2* TSS and the mESC-specific super-enhancer. In sum, our data showed that SnapHiC-D can accurately identify mESC-specific DCC at the *Sox2* locus while alternative methods developed for bulk Hi-C data failed to do so.

### 2.4 SnapHiC-D performance is robust to different input cell numbers

We further evaluated the performance of SnapHiC-D with different numbers of input cells. From the sn-m3c-seq data generated from human brain frontal cortex^9^, we analyzed 1,038 oligodendrocytes (ODC) and 323 microglia (MG), all of which contain more than 150,000 contacts per cell. We fixed the number of cells (N=323) for MG, and performed differential analysis against randomly selected 100, 200, 300,… 900 ODCs using SnapHiC-D. We also included the full list of 1,038 ODCs in the differential analysis.

To benchmark the SnapHiC-D-identified DCCs, we used the H3K4me3 PLAC-seq data generated from ODC and MG^30^. We applied the MAPS pipeline^31^ and identified 21,422 and 41,941 10Kb resolution significant interactions for ODC and MG, respectively. We further defined 12,277 ODC-specific interactions and 32,796 MG-specific interactions, and combined these two sets of cell-type-specific interactions as the reference list. Similar to the previous analysis, we defined a SnapHiC-D-identified DCC as “testable” if it overlaps with a celltype-specific interaction identified from PLAC-seq data. **Figure S3A** and **Figure S3B** showed the number of DCCs and testable DCCs identified using a different number of ODCs, respectively. We observed that a higher number of ODCs led to more ODC-specific DCCs while the number of MG-specific DCCs showed a less pronounced increase. The precision and recall of MG-specific DCCs were robust against different numbers of ODCs (**Figure S3C**, **S3D**), not surprisingly since the number of MG is fixed (N=323). In contrast, the increased number of ODC-specific DCCs resulted in slightly decreasing precision and slightly increasing recall, and both two curves reached a plateau with more than 300 ODCs (**Figure S3C**, **S3D**). Taken together, our data suggest that the performance of SnapHiC-D is robust to different input cell numbers.

### 2.5 DCCs are correlated with the dynamics of gene expression and epigenetics features

Next, we evaluated whether SnapHiC-D-identified DCCs are associated with the dynamic gene expression and epigenetics features. We re-analyzed sn-m3c-seq data generated from mouse hippocampus tissue^32^, where gene expression, chromatin accessibility and five types of histone modification (H3K4me3, H3K4me1, H3K27ac, H3K9me3 and H3K27me3) data of matched cell types in mouse hippocampus are released by our recent Paired-Tag study^33^. We matched the major cell types identified from sn-m3c-seq data and Paired-Tag data and selected three cell types (hippocampal CA1 pyramidal neurons [CA1]: N=408, dentate gyrus [DG]: N=1,040 and oligodendrocytes [ODC]: N=236) with sufficient cell number and sequencing depth (>150,000 contacts per cell) for the downstream differential analysis (**Table S2**).

We first compared the 408 CA1 with the 236 ODC. Among all 744,890 SnapHiC-D-evaluated bin pairs, SnapHiC-D identified 406,552 DCCs (54.6%), where 202,596 and 203,956 DCCs showed higher interaction frequency in CA1 and ODC, respectively. The large proportion of DCCs indicates distinct 3D genome organization between neurons and non-neurons. We further focused on 76,464 DCCs where only one end contains the TSS of one expressed gene (i.e., RPKM>1 in CA1 or ODC), and termed the 10Kb bin containing TSS(s) as the “anchor” bin, and the 10Kb bin not containing TSS as the “target” bin. We found that the significance of DCCs, measured by T-test statistics, is positively correlated with the change of expression of genes within the anchor bins (**Figure 3A**), and the change of the H3K4me3 active promoter mark at the anchor bins (**Figure 3B**). In addition, the significance of DCCs is positively correlated with the change of two active enhancer marks H3K4me1 and H3K27ac, as well as chromatin accessibility at the target bins (**Figure 3C, 3D** and **3E**), and negatively correlated with the change of heterochromatin mark H3K27me3 at the target bins (**Figure 3F**). Interestingly, we observed a weak positive correlation between the dynamic chromatin contacts and the dynamics of H3K9me3 at the target bins (**Figure 3G**). As one illustrative example, we found that CA1-specifically expressed gene *Slc17a7* has 10 DCCs, with interacting target bins containing CA1-specific H3K27ac ChIP-seq peaks (**Figure 4**).

**Figure 3.**
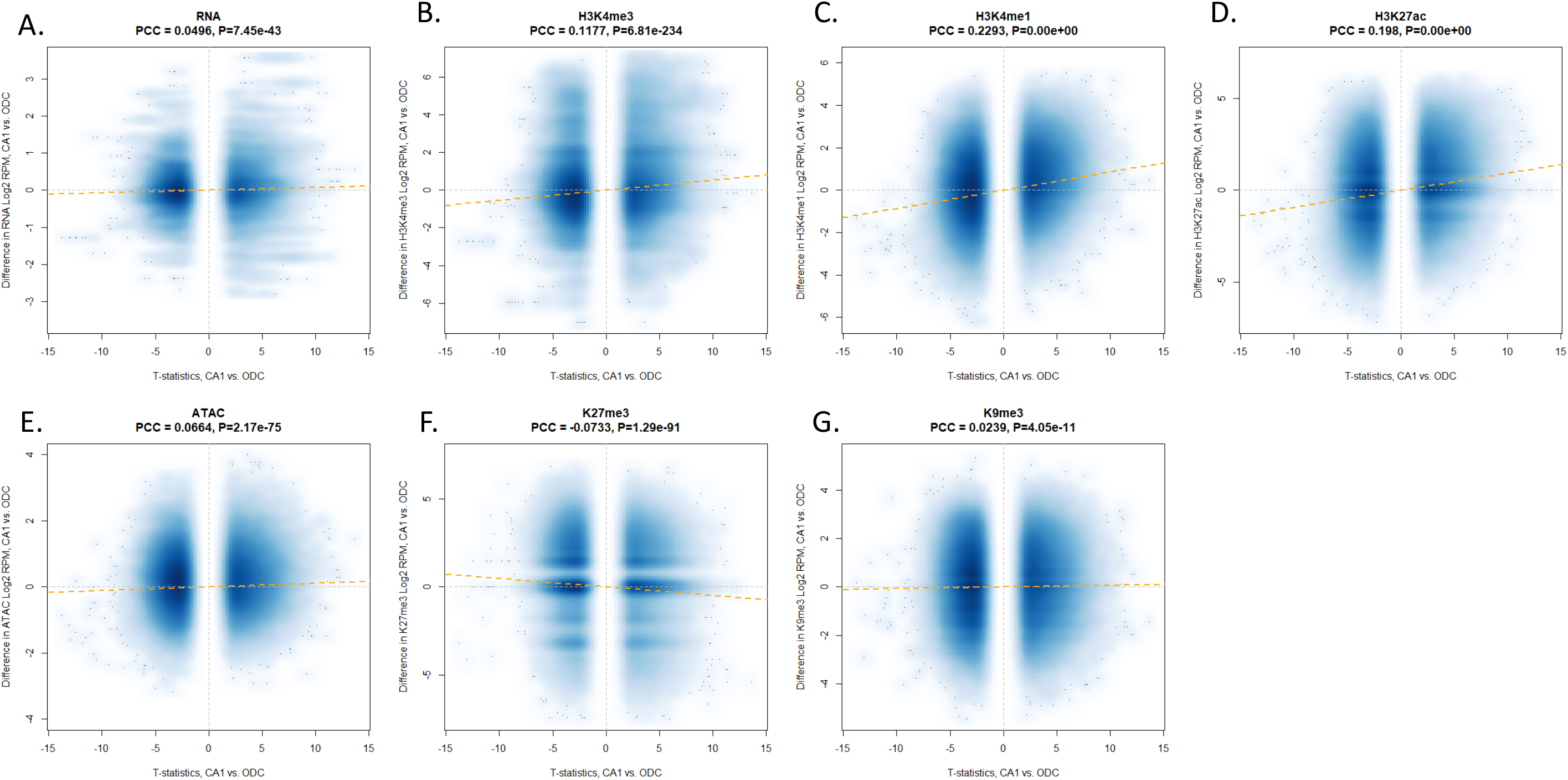
DCCs are correlated with differential gene expression and differential histone marks in mouse hippocampus tissue (408 CA1 vs. 236 ODC). Scatterplot of the significance of DCCs, measured by T-test statistics obtained in SnapHiC-D (x-axis), and the change of gene expression (**A**) and H3K4me3 (**B**) in the anchor bin, and the change of H3K4me1 (**C**), H3K27ac (**D**), ATAC (**E**), H3K27me3 (**F**) and H3K9me3 (**G**) in the target bin. The dashed yellow line is the fitted line between T-test statistics (x-axis) and the change of gene expression or histone marks (y-axis).

**Figure 4.**
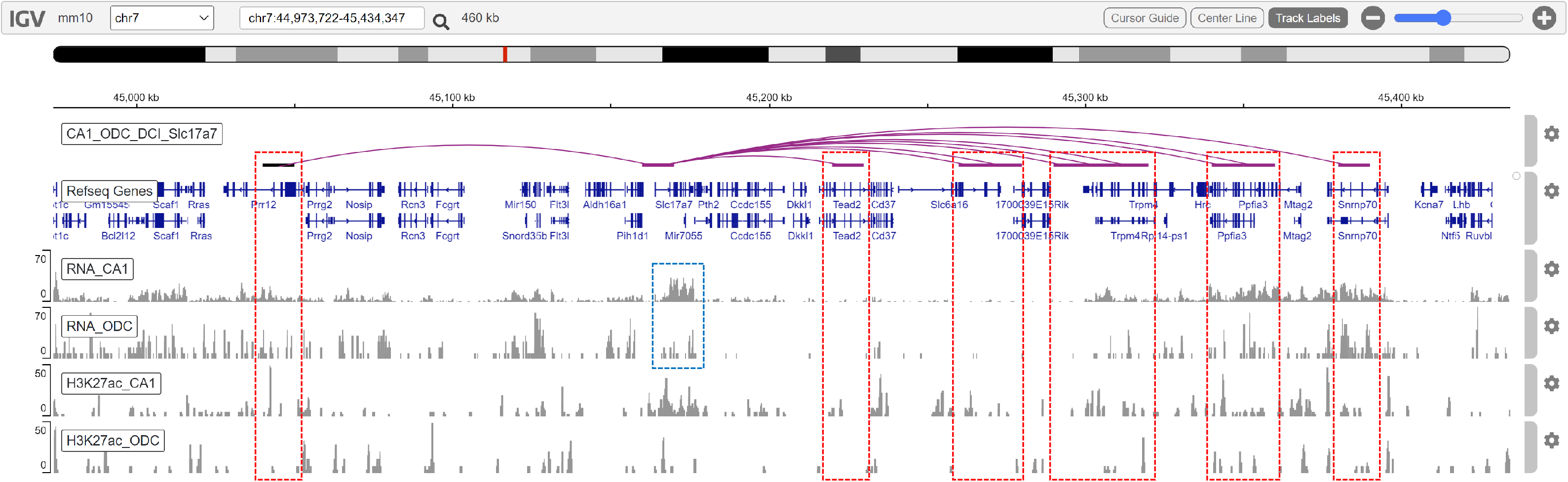
CA1-specifically expressed gene *Slc17a7* has CA1-specific interactions and CA1-specific H3K27ac ChIP-seq peaks. Gene *Slc17a7* is highly expressed in CA1 than ODC, highlighted by the dashed blue box. SnapHiC-D identified 10 DCCs with chromatin interaction frequency higher in CA1 than ODC. The interacting target bins, highlighted by the dashed red boxes, have higher H3K27ac ChIP-seq peaks in CA1 than ODC.

Next, we compared two types of excitatory neurons, CA1 and DG, with 408 and 1,040 cells measured, respectively in the sn-m3c-seq dataset. Among all 606,191 SnapHiC-D-evaluated bin pairs, SnapHiC-D identified 111,404 DCCs (18.4%), where 47,243 and 64,161 DCCs showed higher interaction frequency in CA1 and DG, respectively. Compared to the differential analysis between CA1 and ODC, we observed a smaller proportion of DCCs between CA1 and DG, suggesting that these two types of excitatory neurons share a similar 3D genome. We further focused on 19,391 DCCs where only one end contains the TSS of one expressed gene (i.e., RPKM>1 in CA1 or DG) for downstream correlation analysis. We found that the significance of DCCs is not correlated with the change of expression of genes within the anchor bins (**Figure S4A**), probably due to the similarity of gene expression between CA1 and DG (**Figure S5**). Correlation analysis between 3D genome and epigenome demonstrated consistent results with differential analysis between CA1 and ODC. Specifically, we found that the significance of DCCs is positively correlated with the change of active promoter mark H3K4me3 at the anchor bins (**Figure S4B**), and the change of two active enhancer marks H3K4me1 and H3K27ac, as well as chromatin accessibility at the target bins (**Figure S4C, S4D** and **S4E**), and negatively correlated with change of two heterochromatin marks H3K27me3 and H3K9me3 at the target bins (**Figure S4F**, **S4G**). As one illustrative example, we found that CA1-specificially expressed gene *Kcnq5* has 11 DCCs, with interacting target bins containing CA1-specific H3K27ac ChIP-seq peaks (**Figure S6**).

Additionally, we performed a similar analysis using sn-m3c-seq data generated from human prefrontal cortex tissue^9^, and obtained consistent results (**Supplementary Material Section 2** and **Figure S7**, **S8**). Taken together, our results show that DCCs are correlated with the dynamics of gene expression and epigenetic features in the mammalian genome.

## 3. Conclusion and discussion

In this work, we report SnapHiC-D, a DCC caller tailored to comparative analysis of scHi-C data. We benchmarked SnapHiC-D against two existing DCC callers designed for bulk Hi-C data, BandNorm+diffHiC and multiHiCcompare, using both in-house generated data and public data, and demonstrated the superior performance of SnapHiC-D in terms of sensitivity and accuracy. We applied SnapHiC-D to sn-m3C-seq data generated from mouse hippocampal and human prefrontal cortical tissues, and further revealed that DCCs are correlated with dynamic gene expression and epigenetic marks between distinct cell types.

At least three critical preprocessing steps can affect the quality of DCCs identified from scHi-C data. First of all, we analyzed sn-m3c-seq data released in Liu et al study^32^, where the cell types are pre-defined by DNA methylation data. When DNA methylation data is not available, using scHi-C data alone to define cell type is challenging, in particular, distinguishing cell sub-types, if not impossible^34,35^. Inaccurate cell type clustering results can lead to reduced power in DCC detection.

Second, imputation is indispensable in scHi-C data analysis at kilobase resolution. Several algorithms have been proposed to impute scHi-C data^14,17,35^. Following our published SnapHiC^16^, SnapHiC-D uses the random walk with restart algorithm (RWR) to impute 10Kb resolution scHi-C data. Future studies are warranted to fully benchmark the performance of different imputation algorithms, such as Higashi^17^ and Fast-Higashi^35^, in terms of their impact on DCC detection.

Moreover, data normalization, including both within-cell-type normalization and between-cell-type normalization, plays an important role in DCC detection, similar to the differential analysis in transcriptomic data and other types of genomic data^36–38^. On the one hand, our recent work SnapHiC^16^ demonstrated that RWR-imputed 10Kb resolution chromatin contact probabilities contain negligible systemic biases from restriction enzyme cutting, GC content and sequence uniqueness^39^, therefore the within-cell-type normalization is of less concern. On the other hand, between-cell-type normalization has been extensively studied in DCC analysis of bulk Hi-C data^21,22^, largely due to the cell-type-specific dependency between 1D genomic distance and averaged chromatin contact frequency. SnapHiC-D uses normalized (i.e., Z-score transformed) chromatin contact probabilities as the input for DCC analysis, which implies the same (i.e., 0) average normalized chromatin contact frequency at each 1D genomic distance across different cell types. One caveat is that such between-cell-type normalization may remove cell-type-specific 3D genome features at the global scale^28^. Users need to be cautious with such Z-score transformation when the global scale dynamics of 3D genome are biologically meaningful.

How to evaluate and interpret the identified DCCs is another direction for future research. In this work, our benchmark analysis only used a small proportion of “testable” DCCs among all identified DCCs, due to the lack of a genome-wide gold standard reference DCC list. Data generated from orthogonal technologies, such as SPRITE^40,41^, GAM^42,43^ and super-resolution imaging^44–47^, may serve as a better DCC reference list to benchmark the performance of different DCC callers. In addition, SnapHiC-D focuses on TSS-anchored DCCs to achieve a balance between biological importance and computational efficiency, since the genome-wide search of DCC from scHi-C data requires large computing resources and lacks direct biological interpretation. It is straightforward to modify the search space in SnapHiC-D to bin pairs anchored at *cis*-regulatory elements or loci identified from genome-wide association studies (GWAS). Last but not least, we found a moderate correlation between DCCs and dynamics of histone marks, and a weak correlation between DCCs and dynamics of gene expression. It is well known that gene expression is regulated by multiple levels of epigenetic factors, and the 3D genome is just one type of such factors^48^. Functional perturbation experiments, including MPRA^49^, STARR-seq^50^, CRISPR/Cas9, CRISPRi and CRISPRa are necessary to thoroughly interrogate the molecular functions of dynamic 3D genome on cell-type-specific gene regulation.

This work has focused squarely on DCC detection between different cell types. The methodology can be readily applied to data under different experimental conditions, or data from different groups of individuals (e.g., individuals affected with a certain disease of interest versus individuals not affected). Such data and methods^51–55^ have been burgeoning for other omics data, particularly single cell or single nucleus RNA-seq data. We anticipate similar scHi-C data to be generated where SnapHiC-D will also be valuable.

In sum, we developed SnapHiC-D, a computational pipeline to identify DCCs from scHi-C data. SnapHiC-D has the potential to facilitate a better understanding of chromatin spatial organization in complex tissues and revealing gene regulation mechanisms in a cell-type-specific manner.

## Data availability

Single cell Hi-C data from 94 mESCs and 188 NPCs have been deposited to GEO with accession number GSE210585.

## Acknowledgements

We thank 4D Nucleome consortium investigators for comments and suggestions on the early version of this work. This study was funded by the NIH grants R35HG011922 (to M.H.), UM1HG011585 (to B.R.), U01DA052713 (to Y.L.), and K99HG011483 (to C.Z.). X.L. and Y.L. were also partially funded by the NIH grants R01MH125236, R01GM105785, U24AR076730 and P50HD103573.

## Declaration of competing interest

B.R. is a cofounder and shareholder of Arima Genomics, Inc. and Epigenome Technologies, Inc. The remaining authors declare no competing interest.

## Reference genomes

We used mm10 and hg19 for mouse scHi-C data and human scHi-C data, respectively.

---

**Figure S1.**
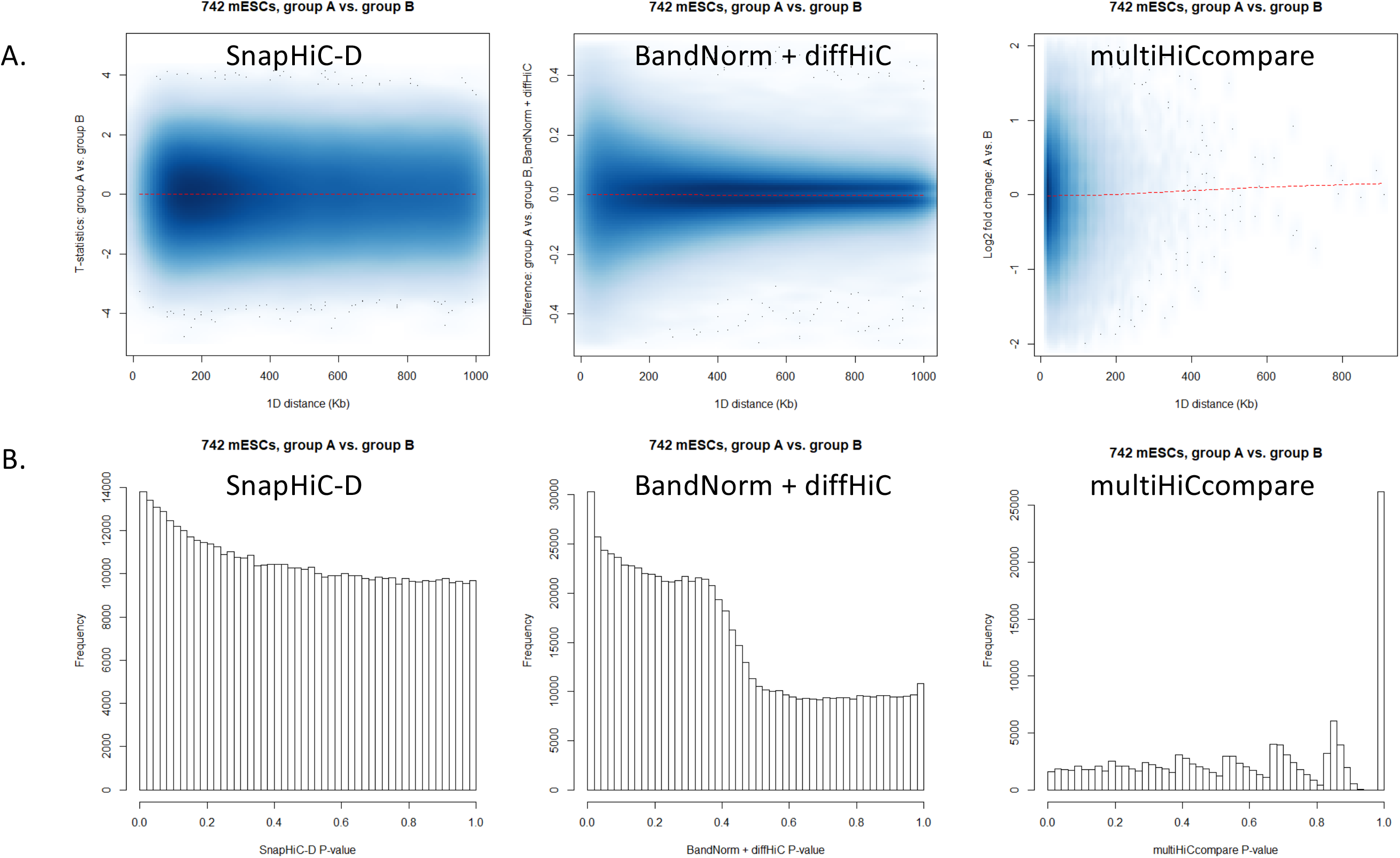
SnapHiC-D controls false positives under the null hypothesis. **A.** MD-plot for SnapHiC-D (left), BandNorm+diffHiC (middle) and multiHiCcompare (right). The dashed red line is the fitting loess curve between 1D genomic distance (x-axis) and the difference in chromatin contact frequency between groups A and B (y-axis). **B.** P-value distribution for SnapHiC-D (left), BandNorm+diffHiC (middle) and multiHiCcompare (right).

**Figure S2.**
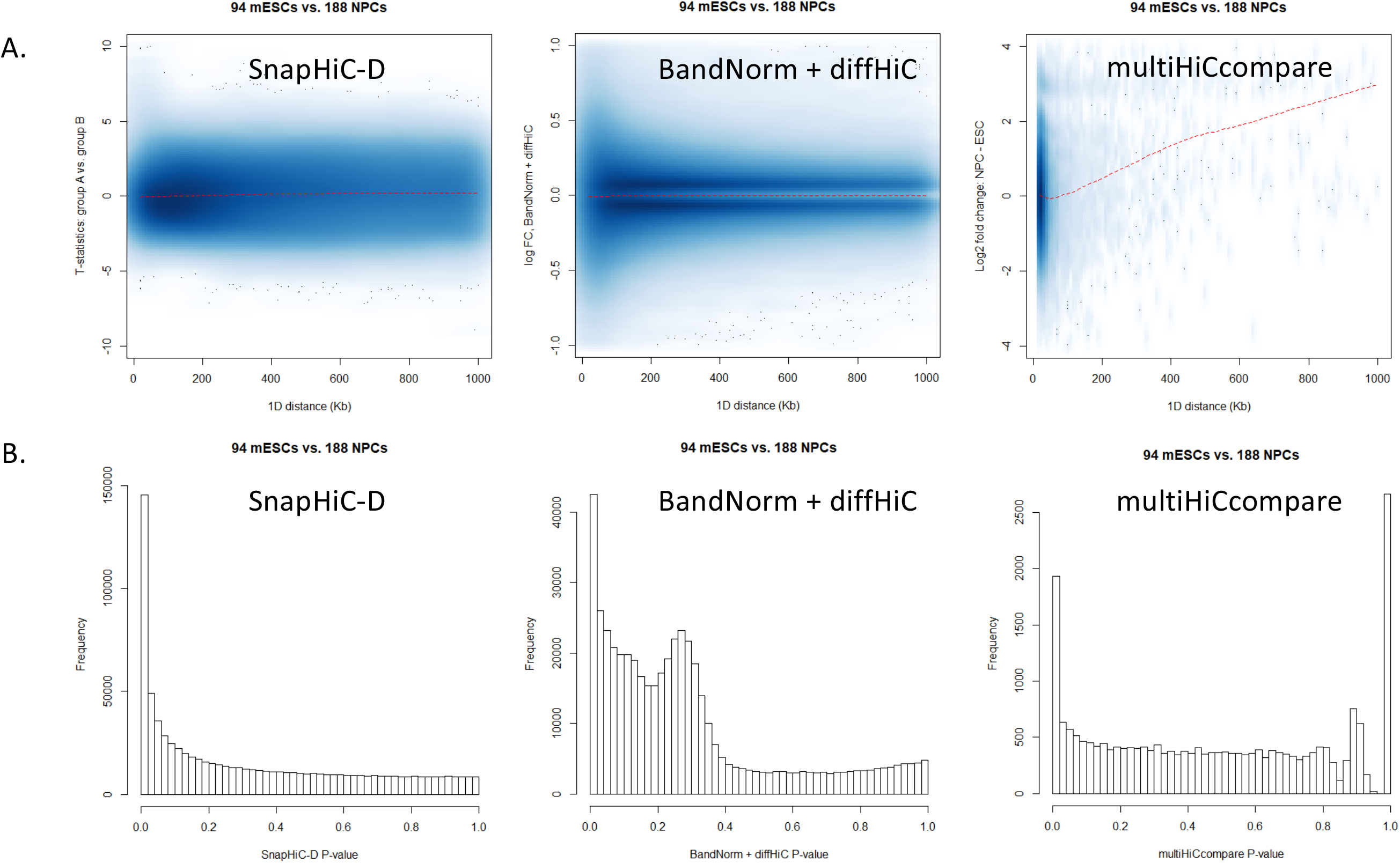
Benchmark the performance of SnapHiC against existing methods using 94 mESCs and 188 NPCs. **A.** MD-plot for SnapHiC-D (left), BandNorm+diffHiC (middle) and multiHiCcompare (right). The dashed red line is the fitting loess curve between 1D genomic distance (x-axis) and the difference in chromatin contact frequency between groups A and B (y-axis). **B.** P-value distribution for SnapHiC-D (left), BandNorm+diffHiC (middle) and multiHiCcompare (right).

**Figure S3.**
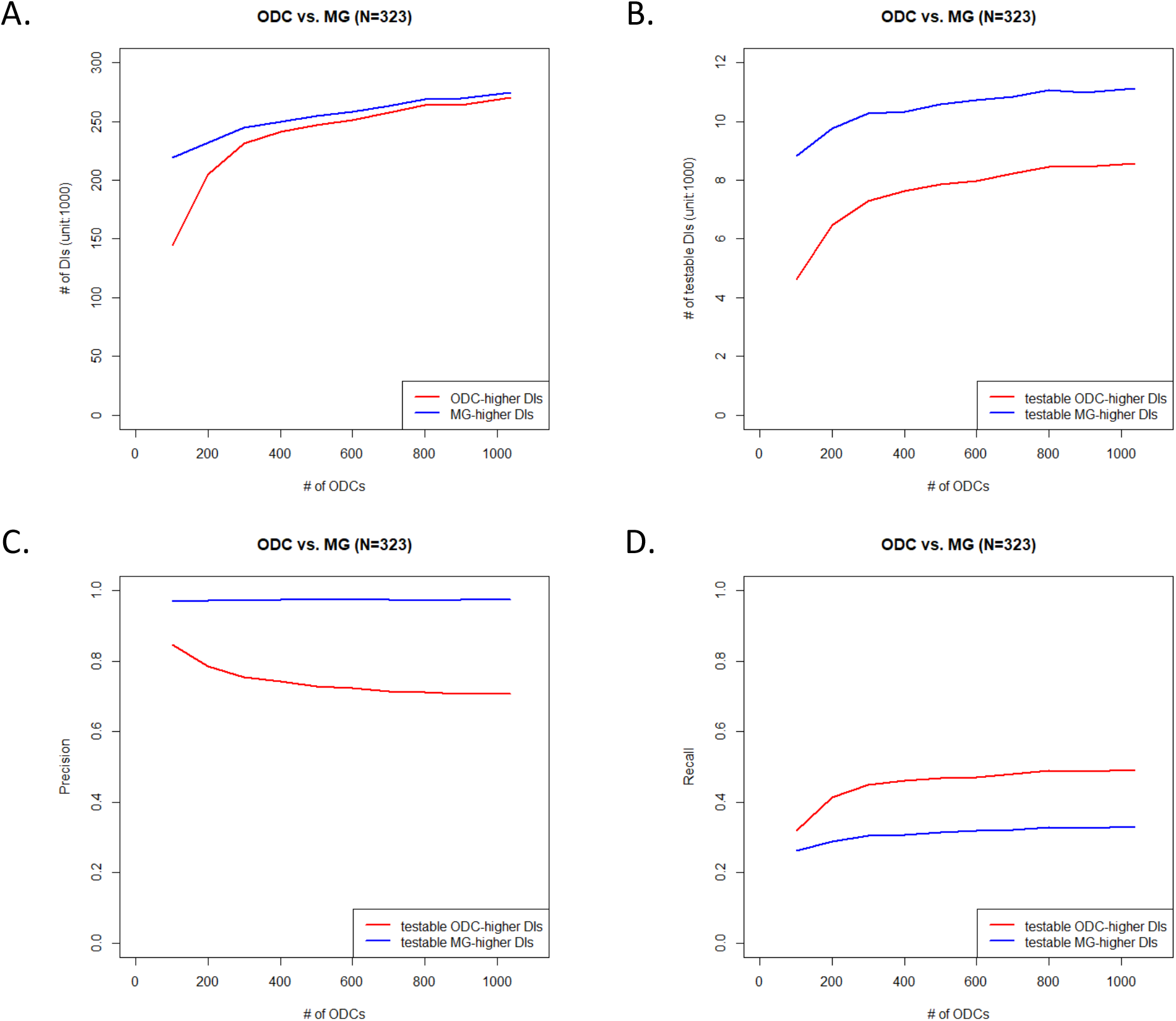
SnapHiC-D performance is robust to different numbers of cells. The number of DCCs (y-axis in **A**) and testable DCCs (y-axis in **B**), when comparing different numbers of ODCs (x-axis) with 323 MG. The precision (y-axis in **C**) and recall (y-axis in **D**), when comparing different numbers of ODCs (x-axis) with 323 MG, using cell-type-specific chromatin interactions identified from H3K4me3 PLAC-seq data as the working truth.

**Figure S4.**
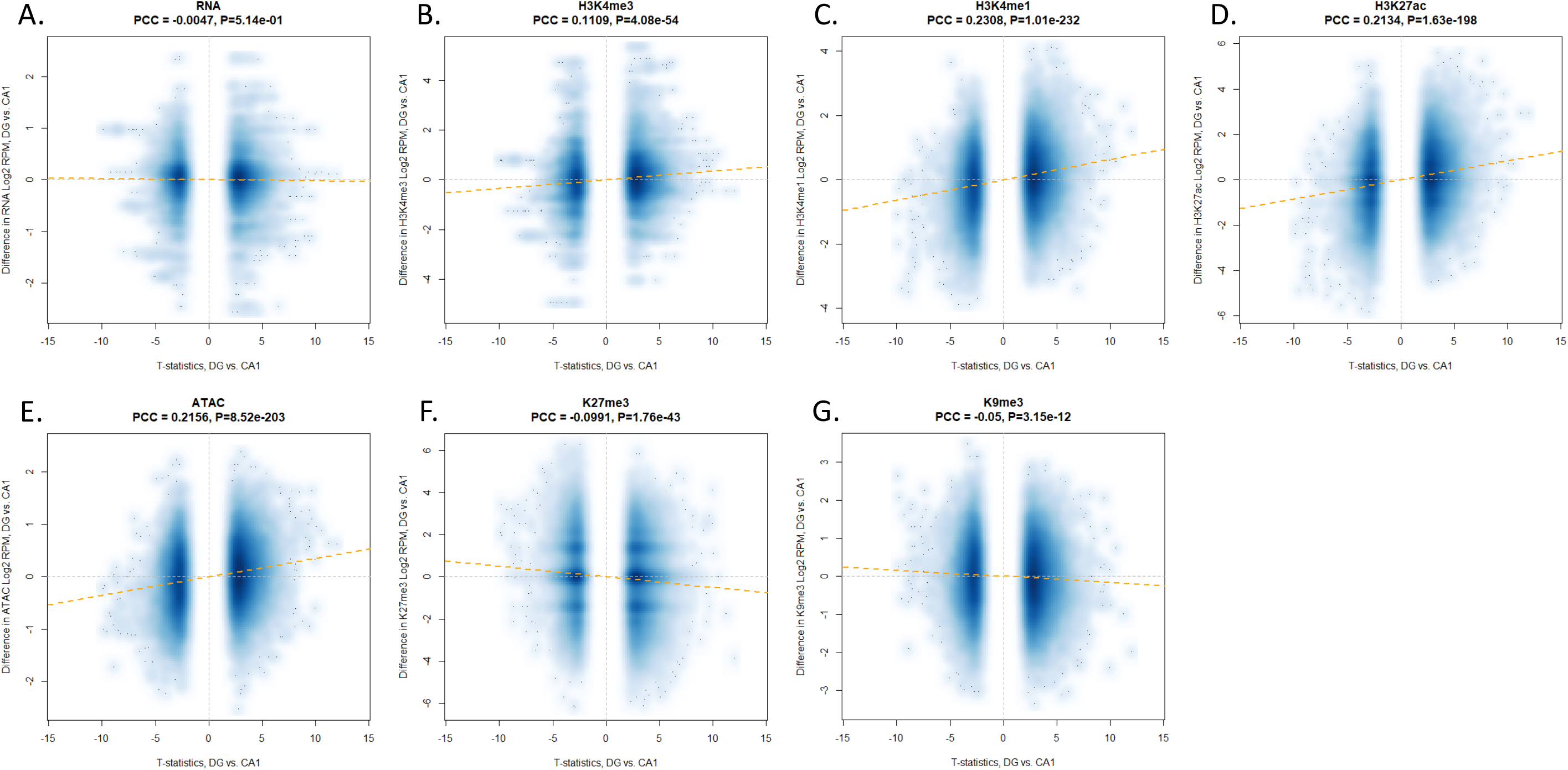
DCCs are correlated with differential histone marks in mouse hippocampus tissue (1,040 DG vs. 408 CA1). Scatterplot of the significance of DCCs, measured by T-test statistics obtained in SnapHiC-D (x-axis), and the change of gene expression (**A**) and H3K4me3 (**B**) in the anchor bin, and the change of H3K4me1 (**C**), H3K27ac (**D**), ATAC (**E**), H3K27me3 (**F**) and H3K9me3 (**G**) in the target bin. The dashed yellow line is the fitted line between T-test statistics (x-axis) and the change of gene expression or histone marks (y-axis).

**Figure S5.**
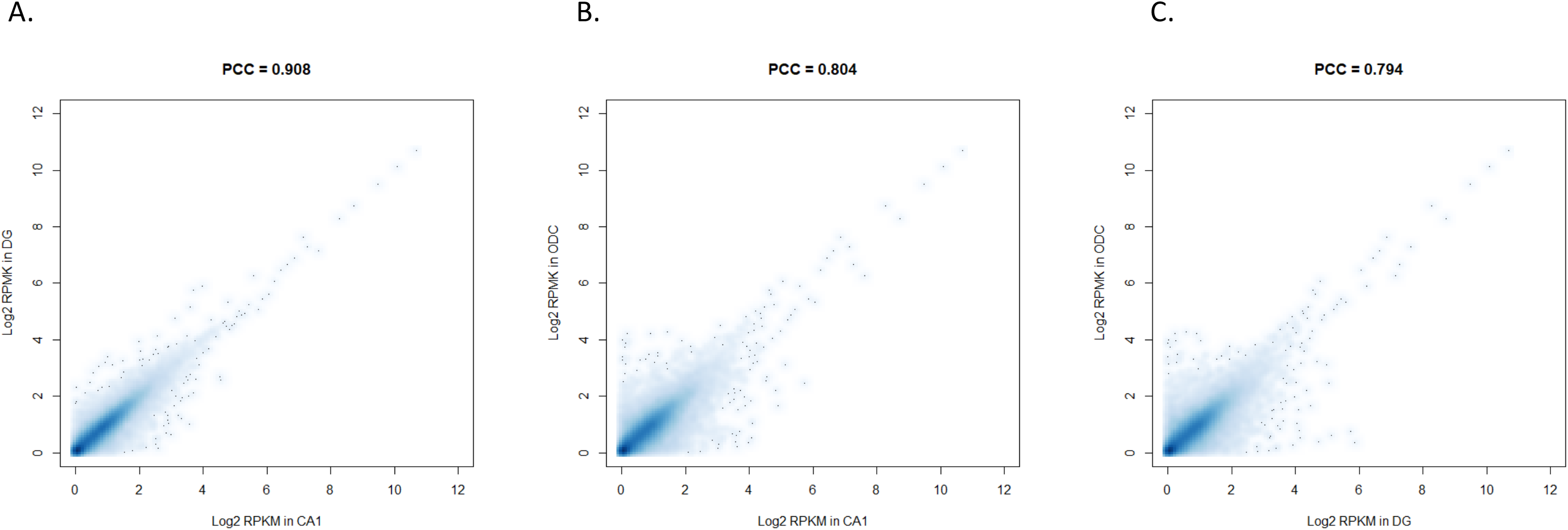
Scatter plot of gene expression data (measured by Log2 RPKM) among CA1, DG and ODC. **(A)** CA1 vs. DG, **(B)** CA1 vs. ODC and (**C**) DG vs. ODC.

**Figure S6.**
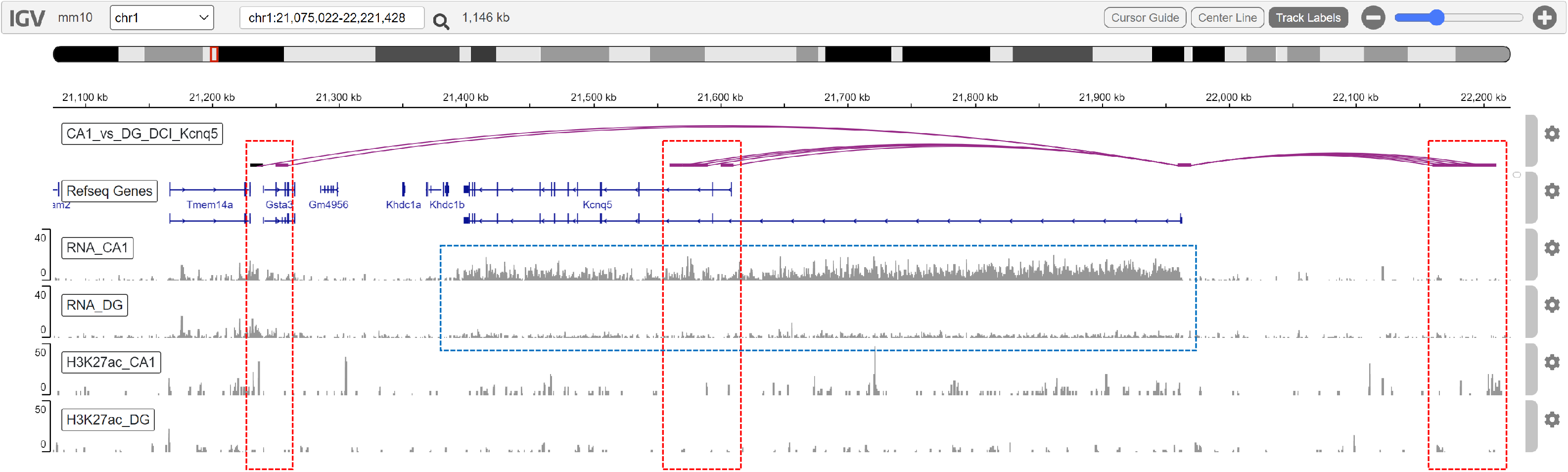
CA1-specifically expressed gene *Kcnq5* has CA1-specific interactions and CA1-specific H3K27ac ChIP-seq peaks. Gene *Kcnq5* is highly expressed in CA1 than DG, highlighted by the dashed blue box. SnapHiC-D identified 11 DCCs with chromatin interaction frequency higher in CA1 than DG. The interacting target bins, highlighted by the dashed red boxes, have higher H3K27ac ChIP-seq peaks in CA1 than DG.

**Figure S7.**
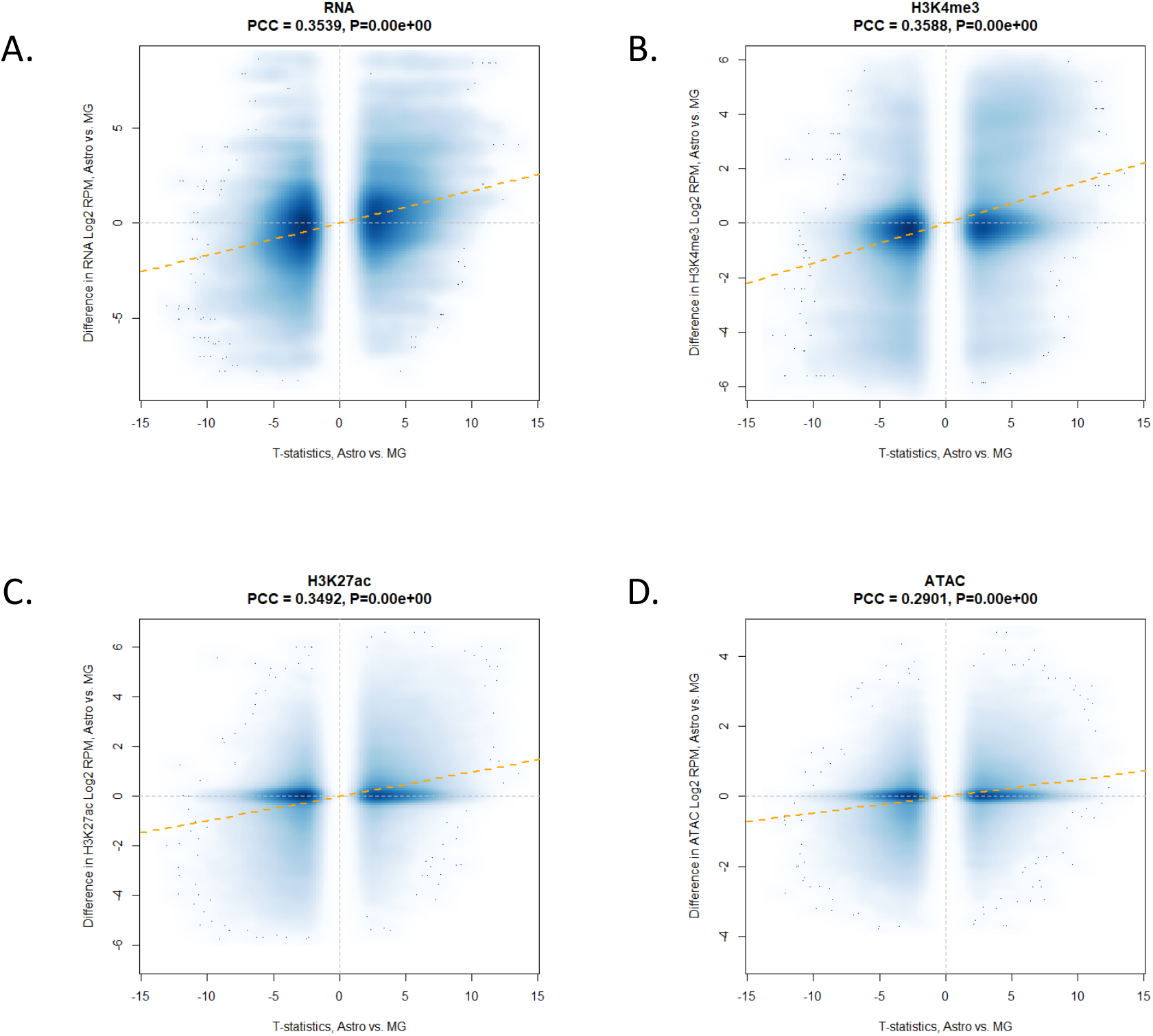
DCCs are correlated with differential gene expression and differential histone marks in human prefrontal cortex tissue (338 Astro vs. 323 MG). Scatterplot of the significance of DCCs, measured by T-test statistics obtained in SnapHiC-D (x-axis), and the change of gene expression (**A**) and H3K4me3 (**B**) in the anchor bin, and the change of H3K27ac (**C**) and ATAC (**D**) in the target bin. The dashed yellow line is the fitted line between T-test statistics (x-axis) and the change of gene expression or histone marks (y-axis).

**Figure S8.**
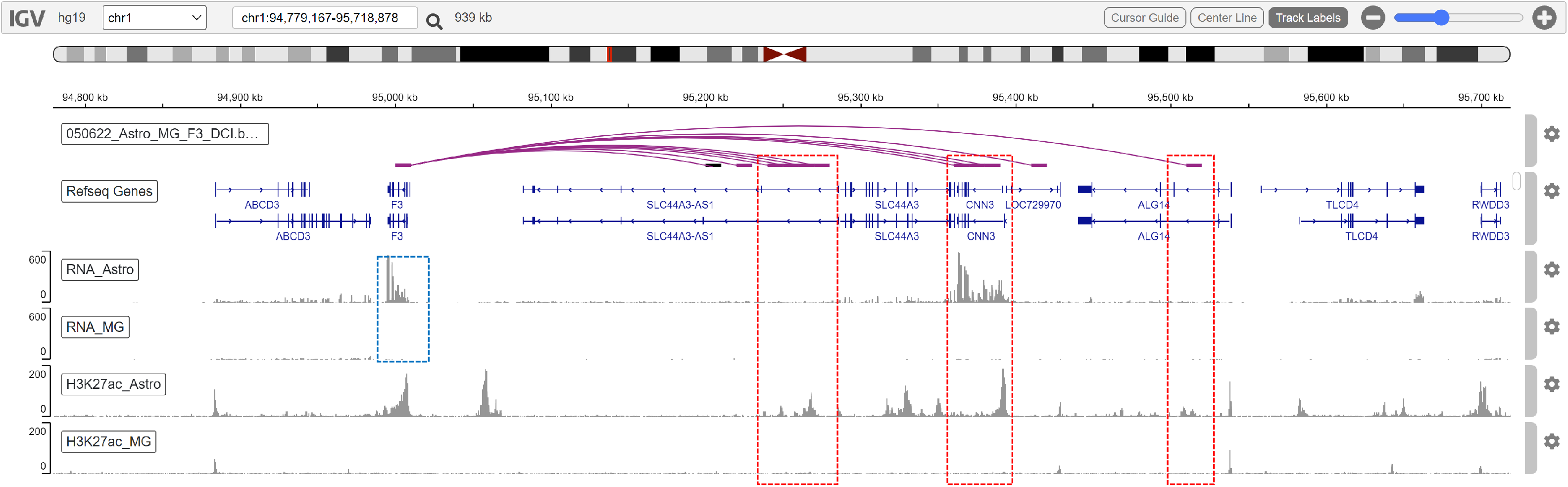
Astro-specifically expressed gene *F3* has Astro-specific interactions and Astro-specific H3K27ac ChIP-seq peaks. Gene *F3* is highly expressed in Astro than MG, highlighted by the dashed blue box. SnapHiC-D identified 11 DCCs with chromatin interaction frequency higher in Astro than MG. The interacting target bins, highlighted by the dashed red boxes, have higher H3K27ac ChIP-seq peaks in Astro than MG.

**Figure S9.**
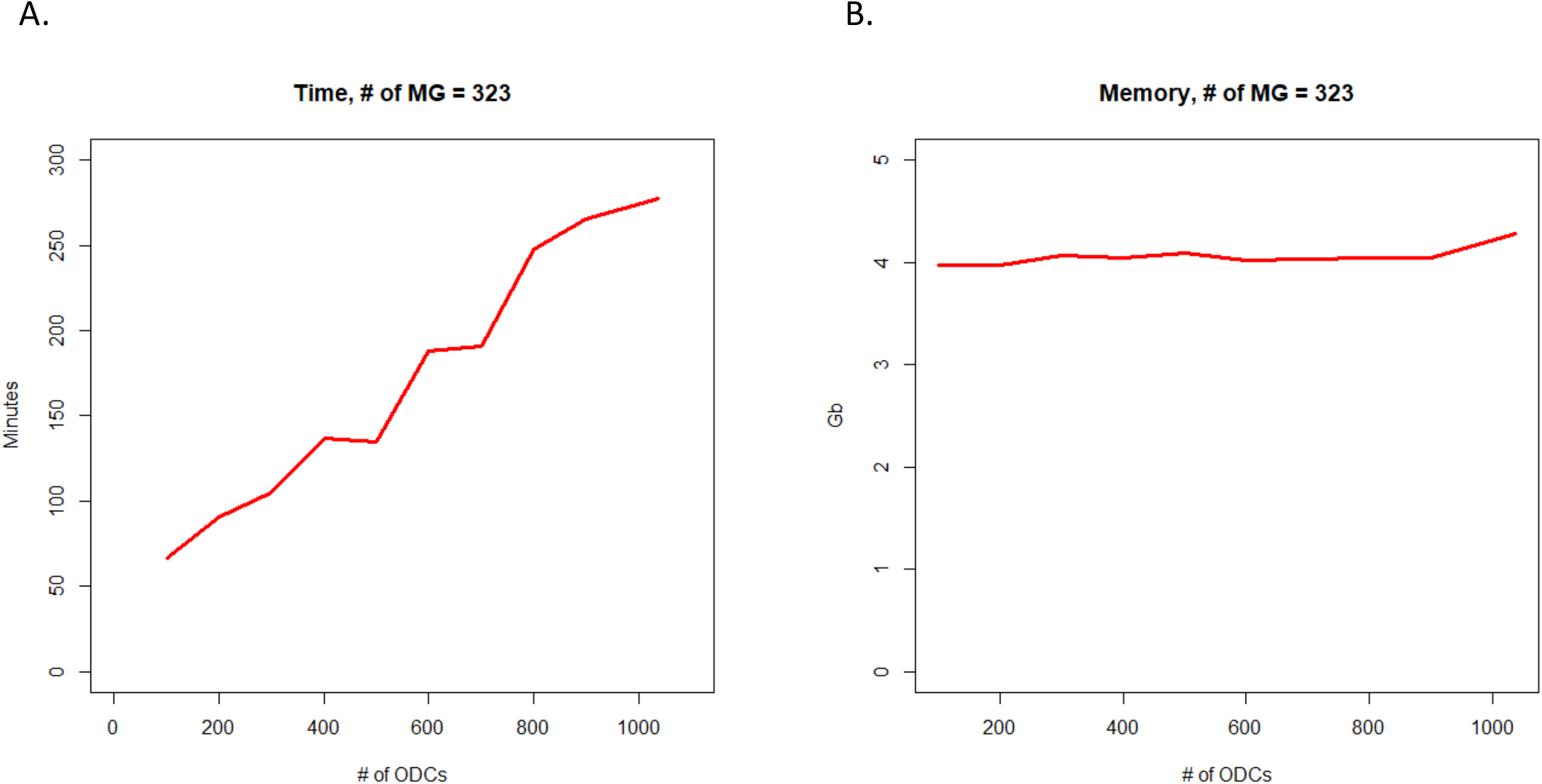
Time cost (A) and memory requirement (B) for SnapHiC-D, comparing different numbers of ODCs with 323 MG.

## Table titles

**Table S1.** Stratified splitting strategy to generate the null hypothesis using 742 mESCs. For all 742 mESCs with >150,000 contacts per cell, the cell cycle information was obtained from the original Nagano et al study^8^. We randomly split cells within the same cell cycle into group A and group B, to account for the potential cell cycle effect on DCCs.

| **Cell cycle** | **Total** | **A** | **B** |
| --- | --- | --- | --- |
| early-S | 301 | 151 | 150 |
| G1 | 129 | 64 | 65 |
| late-S/G2 | 265 | 133 | 132 |
| NA | 29 | 14 | 15 |
| post-M | 4 | 2 | 2 |
| pre-M | 14 | 7 | 7 |
| Total | 742 | 371 | 371 |

**Table S2.** Matched major cell types between sn-m3c-seq data and Paired-Tag data. We matched major cell types defined by sn-m3c-seq data^32^ and Paired-Tag data^33^, both from mouse hippocampus tissue. We then selected three cell types with >150,000 contacts per cell in sn-m3c-seq data for downstream analysis.

|  | Paired-Tag cell type | snm3C-seq Major Type | # of cells with >150K contacts in snm3C-seq |
| --- | --- | --- | --- |
| **Excitatory neurons** | HC CA1/HC subiculum | CA1 | 408 |
|  | HC CA2/3 | CA3/IG-CA2/CA3-St18 | 160 |
|  | HC DG | DG/DG-po | 1,040 |
| **Inhibitory neurons** | InNeu-CGE | CGE-Lamp5/CGE-VIP | 9 |
|  | InNeu-Pvalb | MGE-Pvalb | 6 |
|  | InNeu-Sst | MGE-Sst | 41 |
| **Non-neurons** | OPC | OPC | 36 |
|  | Oligo-MOL/Oligo-MFOL | ODC | 236 |
|  | Endothelial | VLMC/VLMC-pia | 3 |
|  | Microglia | MGC | 83 |
|  | Astro-Nnat/Astro-Myoc | ASC | 37 |

## Methods

### Cell cuture

The *F1 Mus musculus castaneus* × S129/SvJae mouse ESC (F123) line was a gift from Dr. Rudolf Jaenisch and was cultured in KSR medium as previously described^56^. The detailed protocol can be found at 4DN data portal at: https://data.4dnucleome.org/protocols/1d39b581-9200-4494-8b24-3dc77d595bbb/.

To differentiate F123 mESCs from to mNPCs, we followed the previously published protocol^57^. In brief, the F123 mESCs were first passaged once on 0.1% gelatin-coated feeder-free plates before the formation of cellular aggregates (CAs). Then mESCs cells were rthen trypsinized and 4 million cells were plated onto Greiner Petri dishes in CA medium. After four days retinoic acid (RA) was added to CA medium at a final concentration 5μM and the CAs were cultured in CA medium with RA for another four days. After that, the CAs were dissociated and the cells were plated on PORN/laminin-coated plates to allow differentiation of neuronal precursors in N2 medium for another two days before the collection of NPCs.

### Generating scHi-C data

F123 mESCs and NPCs were harvested by accutase treatment and fixed with 1% (v/v) methanol-free formaldehyde at room temperature for 10 minutes. scHi-C libraries were prepared using methods as previously described with slight modifications^58^. In brief, 1-3 million crosslinked mESCs or NPCs were incubated overnight at 37 °C with 200 U MboI followed by proximity ligation at room temperature with slow rotation for 4 hours. Then the nuclei were stained with Hoechst and the single 2N nuclei were sorted by FACS into wells of 96-well plate. After overnight reverse crosslinking at 65 °C, the 3C-ligated DNA in each cell was amplified using GenomiPhi v2 DNA amplification kit (GE Healthcare) for 4-4.5 hours. After purification with AMPure XP magnetic beads and quantification, 10ng WGA product was used to construct library with Tn5.

### scHi-C data preprocessing, imputation and filtering

The fastq files from scHi-C experiments have been mapped and preprocessed following the methods described in the previous studies^8–11^, to generate mapped read pairs (contacts) for each single cell. In brief, the scHi-C read pairs were aligned to mm10 genome with BWA-MEM with the ‘-5’ option, to report the most 5′ end alignment as the primary alignment, and the ‘-P’ option to perform the Smith–Waterman algorithm to rescue chimeric reads. Only the primary alignments were used in the next steps. Then read pairs were de-duplicated with the Picard tool to keep only one read pair at the exact same position. We further filtered the duplicated reads specific to scHi-C datasets: (1) each chromosome was split into consecutive non-overlapping 1-kb bins and only one contact was kept for each 1-kb bin pair, and (2) 1-kb bins that contact with more than ten other 1-kb bins were removed, since they are likely mapping artifacts. Due to the limited ligation capture efficiency, scHi-C data is extremely sparse at kilobase resolution, making the downstream analysis a daunting challenge^18–20^. Different computational approaches, including the random walk with restart algorithm^14^, and the hypergraph-based method Higashi^17^ and Fast-Higashi^35^, have been recently proposed to impute chromatin contact frequency in each cell. Following our recent SnapHiC paper^16^, we also used the RWR imputation in this study. Briefly, we used the RWR algorithm to impute the 10Kb bin resolution contact frequency, for all intra-chromosomal 10Kb bin pairs with the 1D genomic distance between 20Kb and 1Mb, in each cell. Next, we grouped 10Kb bin pairs with the same 1D genomic distance into strata, and normalized the imputed contact frequency into Z-score within each stratum. These Z-scores served as the input file for SnapHiC-D.

We then performed the following three filtering steps to ensure straightforward biological interpretation. First of all, we removed 10Kb bin pairs with either anchor bin overlapping with the ENCODE blacklist regions (http://mitra.stanford.edu/kundaje/akundaje/release/blacklists/mm10-mouse/mm10.blacklist.bed.gz for mm10 and https://www.encodeproject.org/files/ENCFF001TDO/ for hg19) or having low mappability score (<=0.8). The mappability score for each 10Kb bin is defined by our previous study^39^, and can be downloaded from http://enhancer.sdsc.edu/yunjiang/resources/genomic_features/.

Next, we only analyzed 10Kb bin pairs where at least one anchor bin contains transcription start site(s) of protein-coding genes, which are defined by the UCSC refGene annotation (https://hgdownload.soe.ucsc.edu/goldenPath/mm10/bigZips/genes/mm10.refGene.gtf.gz for mm10, and https://hgdownload.soe.ucsc.edu/goldenPath/hg19/bigZips/genes/hg19.refGene.gtf.gz for hg19).

Last but not least, we only kept 10Kb bin pairs that contain more than 10% of outliers (i.e., Z-score >1.96) in at least one cell type. The rationale is to only evaluate bin pairs with sufficiently high contact frequency, and skip bin pairs which are random collisions in both cell types. Similar filtering step has been widely used in differential gene expression analysis, where one only evaluates genes with sufficient gene expression (e.g., RPKM>1) in at least one cell type.

In sum, the final list of filtered bin pairs for SnapHiC-D contains TSS-anchored intra-chromosomal 10Kb bin pairs with 1D genomic distance 20Kb ~ 1Mb, no overlapping with ENCODE blacklist regions or low mappability regions, and with sufficiently high contact frequency in at least one cell type. To make a fair comparison with SnapHiC-D, we also applied the first two filtering steps to create the list of filtered bin pairs for the other methods (BandNorm+diffHiC and multiHiCcompare). Since filtering based on Z-score is specific to the SnapHiC-D algorithm, we replaced it with filtering steps described in other methods, such as filtering bin pairs based on some minimal threshold of raw Hi-C contact frequency in the aggregated pseudo bulk Hi-C data.

### Preprocessing of Paired-Tag data

Paired-Tag data were processed as previously described^33^. Briefly, cellular barcodes were extracted from Read2 and mapped to a reference of all possible cellular barcodes with no more than 1 mismatch; unmapped reads were discarded and low-coverage cells (<1,000 transcripts and <500 DNA fragments) were excluded from further analysis. Reads were then mapped to mm10 reference genome with bowtie2 (for DNA) and STAR (for RNA, with the UCSC refGene annotation), and PCR duplicates were then removed according to the mapped location, UMI, and cellular barcodes. RNA alignment files were then converted to a cell-to-genes count matrix and single-cell clustering was carried out with the Seurat package^59^. DNA alignment files were then converted to cell-to-bins (5Kb) count matrix; the cell type-to-bins count matrix was generated by aggregating counts from cells of the same cell types grouped by RNA-based clustering.

Gene expression levels (for Paired-Tag) and hypo-CH-methylation at gene bodies (for sn-m3c-seq) of marker genes were used to match the major cell types for these two datasets: including CA1 (*Fibcd1*), CA2/3 (*Cacng5*, *Rerg*), DG (*Prox1*), InNeu-CGE (*Vip*), InNeu-Pvalb (*Pvalb*), InNeu-Sst (*Sst*), OPC (*Pdgfra*), ODC (*Mbp*), VLMC (*Slc6a13*), MGC (*Csf1r* and ASC (*Slc1a2*).

### multiHiCcompare

To generate the input files for multiHiCcompare (https://bioconductor.org/packages/release/bioc/html/multiHiCcompare.html), we split each aggregated pseudo bulk Hi-C data into two groups. The input file contains chromosome ID, region 1, region 2 and the raw contact frequency. We used run mulitHiCcompare with the default parameters at the 10Kb bin resolution.

### BandNorm + diffHiC

We first created 10Kb bin resolution contact matrix for each single cell, and applied BandNorm with the default parameters to perform across-cell normalization. We then randomly split cells of the same cell type into two groups, and aggregated them into pseudo bulk Hi-C data, as suggested by the Zheng et al study^27^. Next, we used R package diffHiC to perform differential analysis between two cell types, each consisting of two replicates of aggregated pseudo bulk Hi-C data. We did not use the deep learning-based method 3DVI proposed in the same preprint^27^, since BandNorm coupled with diffHiC achieved comparable or superior performance than 3DVI coupled with diffHiC in terms of DCC detection accuracy (see details in Figure S19F in Zheng et al study^27^).

## Notes

### Competing Interest Statement

Bing Ren is a cofounder and shareholder of Arima Genomics, Inc. and Epigenome Technologies, Inc.

