## Supplementary Materials for "SnapHiC-D: a computational pipeline to identify differential chromatin contacts from single cell Hi-C data"

**1. Benchmark analysis using DCCs identified by applying multiHiCcompare to bulk Hi-C data as the working truth**

In addition to HiCCUPS loops, we treated each replicate as an independent unit, and applied multiHiCcompare to bulk Hi-C data to identify DCCs, named as “bulk-Hi-C-DCCs” for short. Again, we only retained TSS-anchored DCCs. The “bulk-Hi-C-DCC” list consists of 21,641 mESC-specific DCCs, and 16,460 NPC-specific DCCs. We then defined a DCC as “testable” if it overlaps with a bulk-Hi-C-DCC, and determined its status (i.e., true positive or false positive) based on the cell-type-specificity of bulk-Hi-C-DCC. Among all 139,161 SnapHiC-D-identified DCCs, 4,536 DCCs are testable. For the 1,813 mESC-specific testable DCCs, precision and recall are 96.0% and 8.0%, respectively. For the 2,723 NPC-specific testable DCCs, precision and recall are 95.4% and 15.8%, respectively (**Figure 2C**, **2D**). In contrast, among all 80,356 BandNorm+diffHiC-identified DCCs, 1,273 DCCs are testable. For the 650 mESC-specific testable DCCs, precision and recall are 42.2% and 1.7%, respectively. For the 623 NPC-specific testable DCCs, precision and recall are 58.7% and 1.7%, respectively (**Figure 2C**, **2D**). Finally, among all 2,881 multiHiCcompare-identified DCCs, only 16 DCCs are testable. For the 6 mESC-specific testable DCCs, precision and recall are 66.7% and 0.02%, respectively. For the 10 NPC-specific testable DCCs, precision and recall are 70.0% and 0.04%, respectively (**Figure 2C**, **2D**). Taken together, our results demonstrated that SnapHiC-D outperformed BandNorm+diffHiC and multiHiCcompare in both precision and recall, using either bulk Hi-C specific loops or bulk Hi-C DCCs as the working truth.

**2. Differential chromatin contact (DCC) analysis of human brain cell types**

We re-analyzed sn-m3c-seq data generated from human prefrontal cortex tissue^1^, and compared 338 astrocytes (Astro) with 323 microglia (MG), all of which contain more than 150,000 contacts per cell. Among all 1,051,219 SnapHiC-D-evaluated bin pairs, SnapHiC-D identified 509,088 DCCs (48.4%), where 254,468 and 254,620 DCCs showed higher interaction frequency in Astro and MG, respectively. We further focused on 129,359 DCCs where only one end contains the TSS of one expressed gene (i.e., RPKM>1 in Astro or MG), and performed the integrative analysis with RNA-seq data and H3K4me3, H3K27ac ChIP-seq data and ATAC-seq data from Astro and MG^2,3^. We found that the significance of DCCs is positively correlated with the change of expression of genes within the anchor bins (**Figure S7A**), the change of H3K4me3 at the anchor bins (**Figure S7B**), and the change of H3K27ac and chromatin accessibility at the target bins (**Figure S7C**, **S7D**). As one illustrative example, we found that Astro-specifically expressed gene *F3* has 11 DCCs, with interacting target bins containing Astro-specific H3K27ac ChIP-seq peaks (**Figure S8**). These results are consistent with the results obtained from mouse hippocampus tissues (CA1 vs. ODC and CA1 vs. DG).

**3. SnapHiC-D computational cost and memory requirement**

We tested the computational efficiency using different numbers of ODCs (100, 200, 300, …, 900 and 1,038) and 323 MG, and found that the time cost increases linearly as the number of ODCs (**Figure S9A**), while the memory requirement keeps almost constant (**Figure S9B**). Specifically, when compareing 100, 500 and 1038 ODCs with 323 MG, SnapHiC-D took 67, 135 and 277 minutes, respectively. The memory costs for such three comparisons are 4.0Gb, 4.1Gb and 4.3Gb, respectively.

**4. Identification and annotation of cell clusters from sc-m3c-seq data**

In this study, we used the cell clusters definition released by the previous two sc-m3c-seq studies^1,4^, where DNA methylome was used to identify distinct cell clusters. Since DNA methylome data is independent of scHi-C data, we can then apply SnapHiC-D to detect DCCs. When DNA methylome data is not available, it is possible to use scHi-C data alone to identify distinct cell clusters^5-7^. However, we advise against applying SnapHiC-D to distinct cell clusters defined by scHi-C data, since it is a circular analysis (i.e., double-dipping problem), which has been well studied in scRNA-seq differential analysis^8^.
